# Viruses are a dominant driver of protein adaptation in mammals

**DOI:** 10.1101/029397

**Authors:** David Enard, Le Cai, Carina Gwenapp, Dmitri A. Petrov

## Abstract

Viruses interact with hundreds to thousands of proteins in mammals, yet adaptation against viruses has only been studied in a few proteins specialized in antiviral defense. Whether adaptation to viruses typically involves only specialized antiviral proteins or affects a broad array of proteins is unknown. Here, we analyze adaptation in ~1,300 virus-interacting proteins manually curated from a set of 9,900 proteins conserved across mammals. We show that viruses (i) use the more evolutionarily constrained proteins from the cellular functions they hijack and that (ii) despite this high constraint, virus-interacting proteins account for a high proportion of all protein adaptation in humans and other mammals. Adaptation is elevated in virus-interacting proteins across all functional categories, including both immune and non-immune functions. Our results demonstrate that viruses are one of the most dominant drivers of evolutionary change across mammalian and human proteomes.

## Introduction

A number of virus-interacting proteins (VIPs) with a specialized role in antiviral defense have been shown to have exceptionally high rates of adaptation (Cagliani et al., 2012; Cagliani et al., 2011; Elde et al., 2009; Fumagalli et al., 2010; Kerns et al., 2008; Liu et al., 2005; Sawyer et al., 2004, 2007; Sawyer et al., 2005; Sironi et al., 2012; Vasseur et al., 2011). One example is protein kinase R (PKR), which recognizes viral double-stranded RNA upon infection, halts translation, and as a result blocks viral replication (Elde et al., 2009). PKR is one of the fastest adaptively evolving proteins in mammals. Specific amino acid changes in PKR have been shown to be associated with an arms race against viral decoys for the control of translation (Elde et al., 2009).

However, PKR and other fast-evolving antiviral defense VIPs are not representative of the hundreds to thousands of other VIPs. Viruses enter and hijack the cells of their hosts by physically interacting with hundreds to thousands of host proteins. Most VIPs are not specialized in antiviral defense, and do not even have any known role in immunity. Many VIPs instead have key functions in basic cellular processes subverted by viruses, and viruses tend to interact with proteins that are functionally important hubs in the protein-protein interaction network of the host (Dyer et al., 2008; Halehalli and Nagarajaram, 2015). It is plausible that many VIPs might evolve to limit the impact of the viruses on the host. However, it is unknown whether the war against viruses is fought by a “professional” army of specifically antiviral proteins, or whether it is a global war fought by a broad range of VIPs.

One reason to believe that the war against viruses might not affect evolution of a broad array of VIPs is that, contrary to the pattern observed for specifically antiviral proteins, most VIPs evolve unusually slowly (Davis et al., 2015; Jager et al., 2012). Furthermore, very few cases of adaptation to viruses have been studied outside of fast evolving, specialized antiviral proteins. Transferrin receptor or TFRC is the most notable exception, as a striking example of a non-immune, housekeeping protein hijacked by viruses (Demogines et al., 2013; Kaelber et al., 2012). TFRC is responsible for iron uptake in many different cell types and is used as a cell surface receptor by diverse viruses in rodents and carnivores. TFRC has repeatedly evaded binding by viruses through recurrent adaptive amino acid changes. As such, it is the only known clear case of a host protein with adaptation in response to viruses that is not specialized in the antiviral response. Whether TFRC represents an exception or the rule for VIPs is currently unknown.

Whether the slower evolution of VIPs primarily reflects lower rates of adaptation, greater evolutionary constraint, or both is currently unknown. Here we analyze patterns of evolutionary constraint and adaptation in a manually curated, high quality set of ~1,300 VIPs. The vast majority of these VIPs (80%) have no known antiviral or any other more broadly defined immune activity. We confirm that VIPs tend to evolve slowly and show that this is because VIPs experience stronger evolutionary constraint than other proteins within the same functional categories. Most importantly, we show that despite this greater evolutionary constraint, VIPs have experienced much higher rates of adaptation in humans and other mammals compared to other proteins. This excess of adaptation is visible in VIPs involved in most biological functions hijacked by viruses (such as transcription or signal transduction), showing that adaptation in response to viruses extends well beyond the usual, specialized antiviral proteins. We demonstrate that viruses have driven a substantial proportion of all adaptations across the human and mammalian proteomes, establishing that the war against viruses does indeed affect the proteome as a whole.

We finally showcase the power of our global scan for adaptation in VIPs by studying the case of aminopeptidase N, a well-known multifunctional enzyme (Mina-Osorio, 2008) used by coronaviruses as a receptor (Delmas et al., 1992; Yeager et al., 1992). Using our approach we reach an amino-acid level understanding of parallel adaptive evolution in aminopeptidase N in response to coronaviruses in a wide range of mammals.

## Results

Here we analyze patterns of both adaptive evolution and evolutionary constraint/purifying selection in a large set of 1,256 manually curated VIPs from the low-throughput virology literature (Methods and Table S1 available online). We exclude interactions identified by high-throughput experiments as we are concerned about a high rate of false positives (Mellacheruvu et al., 2013). These 1,256 VIPs were annotated from an initial set of 9,861 proteins with orthologs in 24 mammals with high quality genomes (Figure S1, Table S2 and Methods). VIPs in our dataset interact with viral proteins, viral RNA, or viral DNA. Most of them (95%) correspond to an interaction between a human protein and a virus infecting humans (Table S1). Human Immunodeficiency Virus type 1 (HIV-1) is the best-represented virus with 240 VIPs, with nine other viruses having at least 50 VIPs (Table S1).

This dataset represents the largest, most up-to-date set of VIPs backed up by individual low-throughput publications. Nonetheless, given that many VIPs were discovered only recently, with half of all publications reporting VIPs published in the past 7 years (Figure S2), it is likely that additional VIPs remain to be discovered.

These 1,256 VIPs are involved in very diverse cellular processes, as shown by their representation in the Gene Ontology (GO) annotation system of cellular biological processes (2015; Ashburner et al., 2000) (Table S3) such as transcription (354 VIPs), post-translational protein modification (224 VIPs), signal transduction (396 VIPs), apoptosis (185 VIPs), and transport (264 VIPs). The supracellular processes notably include defense response (103 VIPs) and developmental processes (327 VIPs). Many VIPs have no known specifically antiviral immune activity (defined here as any activity restricting viral replication, Methods), and only 241 or 20% of VIPs have any known immune function (Methods and Table S4). These 241 immune VIPs include the VIPs classified as antiviral (Table S4) throughout this manuscript. In total, 162 overlapping GO cellular and supracellular processes have more than 50 VIPs (Table S3). These observations confirm that viruses interact with proteins involved in the majority of basic cellular processes.

## Patterns of purifying selection in VIPs

We confirm the observations of several recent studies suggesting that VIPs tend to evolve slowly (Davis et al., 2015; Jager et al., 2012). The VIPs have ~15% lower mammals-wide dN/dS ratio on average compared to non-VIPs (0.124 versus 0.145, 95% CI [0.136,0.148]; Methods). The difference in dN/dS is highly significant as shown by a permutation test where VIPs are compared with the same number of randomly chosen non-VIPs many times (simple permutation test *P*=0 after 10^9^ iterations; Table S2).

To disentangle whether the slower evolution of VIPs is due to stronger purifying selection or to a lower rate of adaptation, we use the ratio of non-synonymous polymorphisms to synonymous polymorphisms pN/pS rather than the dN/dS ratio. Unlike dN/dS that is strongly influenced by both the effects of purifying selection and adaptation, pN/pS is primarily determined by the efficiency of purifying selection in removing deleterious non-synonymous mutations.

Genome-wide polymorphisms required to measure pN/pS at the scale of the proteome have become available for humans (Abecasis et al., 2012) (1,000 Genomes Project) (Table S5), and chimpanzee, gorilla, and orangutans (Prado-Martinez et al., 2013) (Great Apes Genome Project) (Table S6). The 1,000 Genomes Project and the Great Apes Genome Project are complementary for this analysis. On the one hand, the 1,000 Genomes Project provides high quality variants with frequencies estimated from a large number of individuals. On the other hand the Great Ape Genome project includes fewer individuals, but provides substantial pN and pS counts for more genes than the 1000 genomes data. This is because the great apes populations are more polymorphic than modern human populations, and are less affected by the noise due to genetic drift and strong bottlenecks. Thus, although the 1,000 genomes are well suited for approaches requiring precise frequency estimates (such as the McDonald-Kreitman test implemented below (Messer and Petrov, 2013)), the Great Ape genome data is well suited for the estimation of the pN/pS ratio in as many proteins as possible. Specifically, we measure pN/pS as the average across non-human great apes (or as the average in the 1,000 Genomes African populations; Supplemental Methods) using the data from the largest chimpanzee, gorilla, and orangutan populations in order to further limit the noise in the estimation of the strength of purifying selection due to drift and bottlenecks (Prado-Martinez et al., 2013). We use only great ape pN and pS in the analyses that involve subsamples of proteins in order to retain sufficient statistical power.

In human African populations from the 1,000 Genomes project (Supplemental Methods), the average pN/pS is 21% lower in VIPs compared to non-VIPs (0.759 versus 0.966, 95% CI [0.92,1.01], simple permutation test *P*=0 after 10^9^ iterations). This suggests that in the human lineage VIPs have been under stronger purifying selection than non-VIPs. In line with this, VIPs also show an excess of low frequency (≤10%) deleterious non-synonymous variants compared to non-VIPs (Figure S3). In great apes, the average pN/pS ratio is 25% lower in VIPs compared to non-VIPs (0.526 versus 0.697, 95% CI [0.66,0.72], simple permutation test *P*=0 after 10^9^ iterations). The distribution of pN/pS in VIPs is globally skewed towards lower values compared to non-VIPs (Figure 1A). The difference in pN/pS in great apes between VIPs and non-VIPs is robust to a number of potentially confounding factors, including gene length, GC content and recombination (Table S7). These results show that VIPs do experience stronger purifying selection than non-VIPs. Finally, stronger purifying selection acting on the VIPs is widespread and is not limited to VIPs interacting with any one particular virus (Figure 1B).

**Figure 1.**
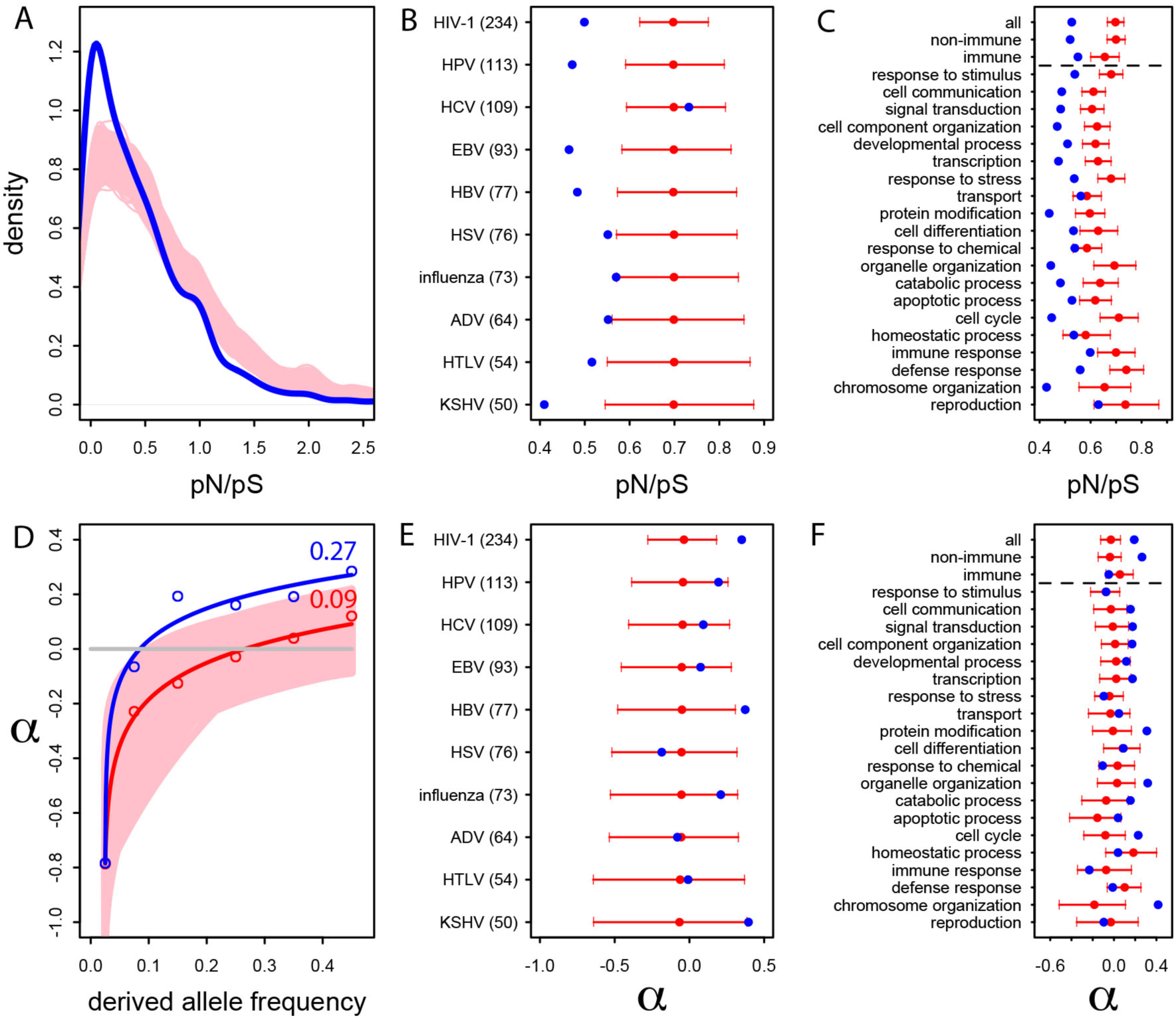
Patterns of purifying selection and human adaptation in VIPs. A) Distribution of pN/pS in VIPs (blue) and non-VIPs (pink). The blue curve is the density curve of pN/(pS+1) for 1,256 VIPs. We use pN/(pS+1) instead of pN/pS to account for those coding sequences where pS=0. pN and pS are measured using great ape genomes from the Great Ape Genome Project (Supplemental Methods). The pink area represents the superimposition of the density curves for each of 5,000 sets of randomly sampled non-VIPs. B) Average pN/pS in VIPs (blue dot) versus average pN/pS in non-VIPs (red dot and red 95% confidence interval) within ten viruses with more than 50 VIPs The number between parentheses is the number of VIPs for each virus. KSHV: Kaposi’s Sarcoma Herpesvirus. HIV-1: Human Immunodeficiency Virus type 1. HBV: Hepatitis B Virus. ADV: Adenovirus. HPV: Human Papillomavirus. HSV: Herpes Simplex Virus. EBV: Epstein-Barr Virus. Influenza: Influenza Virus. HTLV: Human T-lymphotropic Virus. HCV: Hepatitis C virus. C) Same as B), but for the 20 most high level GO processes with the highest number of VIPs. The full GO process name for “protein modification” as written in the figure is “post-translational protein modification”. D) Asymptotic MK test (Supplemental Methods) for the proportion of adaptive amino acid substitutions (a) in VIPs (blue dots and curve) and non-VIPs (red dots and curve). Pink area: superposition of fitted logarithmic curves for 5,000 random sets of 1,256 non-VIPs (as many as VIPs) where the estimated a falls within a’s 95% confidence interval. E) Classic MK test (Supplemental Methods) for VIPs (blue dot) and non-VIPs (red dot and 95% confidence interval) for the ten viruses with 50 or more VIPs. F) Same as E) but for the 20 top high level GO processes with the most VIPs below the dotted black line. Above the dotted black line: the classic MK test for all VIPs, for non-immune VIPs and for immune VIPs (Table S4). See also Tables S3, S4, S5, S6, S7, S8 and S9 and Figures S3 and S4

The higher level of purifying selection in VIPs might be due to the fact that VIPs participate in the more constrained host functions, or, alternatively, because within each specific host function, viruses tend to interact with the more constrained proteins. In order to assess these two non-mutually exclusive scenarios we generated 10^4^ control sets of non-VIPs chosen to be in the same 162 Gene Ontology processes as VIPs (GO processes with more than 50 VIPs; Table S3 and Methods).

In great apes, GO-matched non-VIPs still have a much higher pN/pS ratio compared to VIPs, suggesting that VIPs tend to be more conserved than non-VIPs from the same GO category. On average, pN/pS in the GO-matched non-VIPs is 0.647 (95% CI [0.621,0.674]). This is only slightly lower than the average ratio in non-VIPs in general (pN/pS=0.697, *P*=2×10^−3^), but much higher than the average ratio in VIPs (0.526, permutation test *P*=0 after 10^9^ iterations). The stronger purifying selection acting on VIPs is apparent within most functions. Figure 1C shows stronger purifying selection in the 20 high level GO categories with the most VIPs. In all the 20 GO categories pN/pS is lower in VIPs than in non-VIPs, and the difference is significant for 17 of these categories (Table S3). This shows that within a wide range of host functions, viruses tend to target the most conserved proteins.

Interestingly, even immune VIPs (Table S4) have a significantly reduced pN/pS ratio compared to immune non-VIPs (Figure 1C), which suggests that immune proteins in direct contact with viruses are more constrained. The reduction in pN/pS in non-immune VIPs (no antiviral or any other immune function, Table S4) is very similar to the reduction observed in the entire set of VIPs (Figure 1C). Table S3 further shows stronger purifying selection in 124 of the 162 GO categories (77%) with more than 50 VIPs. These results suggest that viruses target the most constrained proteins within many diverse host functions, rather than specifically targeting only those host functions that are under strong evolutionary constraint.

## Frequent adaptation in VIPs in the human lineage

Our pN/pS analysis demonstrates that viruses tend to interact with proteins that are under greater evolutionarily constraint. However, this does not necessarily imply that VIPs experience less adaptation than non-VIPs. We estimate the proportion of adaptive non-synonymous substitutions (noted *α*) in VIPs and non-VIPs in the human lineage by using the McDonald-Kreitman test (MK test) (Methods). We first use the classic MK test (McDonald and Kreitman, 1991) with the 1,000 Genomes Project polymorphism data from African populations (Supplemental Methods and Table S5). Since VIPs are more constrained than non-VIPs and tend to have more non-synonymous deleterious low frequency variants than non-VIPs (Figures 1 and S3), we limit the effect of deleterious variants by excluding all variants with a derived allele frequency lower than 10% (Supplemental Methods) (Charlesworth and Eyre-Walker, 2008; Eyre-Walker and Keightley, 2009; Keightley and Eyre-Walker, 2007; Messer and Petrov, 2013). We find that *α* is strongly elevated in VIPs compared to non-VIPs (*α*=0.19 in VIPs versus -0.02 in non-VIPs, permutation test *P*=2.×10^−5^). This shows that VIPs have a substantial excess of adaptation compared to non-VIPs.

The classic MK test is known to be biased downward by the presence of slightly deleterious non-synonymous variants and this bias is difficult to eliminate fully even by excluding low frequency variants (Messer and Petrov, 2013). Note that our application of the classic MK test to discover the higher rate of adaptation in VIPs compared to non-VIPs is conservative given that the VIPs have a higher proportion of slightly deleterious non-synonymous variants and thus should show a stronger downward bias in the estimation of *α* (Figure S3). However, the nominal estimates of *α*=0.19 and -0.02 in VIPs and non-VIPs are likely underestimates of the true proportions of adaptive amino acid changes.

We therefore apply an asymptotic modification of the MK test known to provide estimates of *α* without a downward bias in the presence of slightly deleterious variants (Messer and Petrov, 2013). To further validate the asymptotic MK test we carry out extensive population simulations (Messer, 2013) to show that this test is indeed robust to a number of potential biases (Supplemental Methods and Table S8).

Using the asymptotic MK test we estimate that in VIPs, ~27% of the 1,897 amino acid substitutions along the human lineage were adaptive (Figure 1D). This proportion is three times higher than the estimated proportion of ~9% in non-VIPs (Figure 1D). Thus, although VIPs represent only 13% of the orthologs in our dataset, we estimate that in human evolution they account for almost 30% of all adaptive amino-acid changes. Note that both VIPs and non-VIPs in our dataset are limited to the proteins conserved across all mammals.

The high *α* in VIPs is not explained by higher rates of adaptation in the host GO processes where VIPs are well represented (Figure S4 and Methods). Furthermore, the large difference in *α* observed between VIPs and non-VIPs is robust to a number of potentially confounding factors such as recombination, GC content or gene length (Table S9 and Supplemental Methods). The lower pN/pS in VIPs does not explain their higher *α* either (Table S9).

We further use the classic MK test (excluding variants below 10%) to investigate the excess of adaptation for the specific VIPs of ten human viruses and in the 20 high level GO categories with the most VIPs (Figure 1E and F). We do not use the asymptotic MK test because it does not have sufficient precision when using a small number of proteins (Supplemental Methods). Although the small number of proteins interacting with individual viruses precludes precise estimates of *α* (see the large confidence intervals on Figure 1E), the VIPs show nominally higher values of *α* for eight out of 10 viruses, with HIV-1 and Hepatitis B Virus (HBV) displaying statistically significant increases in adaptation. Likewise, VIPs in most GO categories show higher rates of adaptation (14 out of 20) with 9 of 14 showing statistically significant increases (Figure 1F).

Finally and importantly, the 80% of VIPs with no known antiviral or broader immune function (Table S4) have a strongly increased rate of adaptation according to both the classic MK test (*α*=0.26 in VIPs versus -0.02 in non-VIPs, permutation test *P*=3×10^−7^; Figure 1F) and the asymptotic MK test, with the latter estimating *α*=38% in non-immune VIPs against only 11% for non-immune non-VIPs. Intriguingly, unlike for non-immune VIPs or all VIPs considered together (top of Figure 1F), immune VIPs, including antiviral VIPs (Table S4), do not show any increase of adaptation compared to immune non-VIPs. We speculate that this pattern might reflect the masking effect of balancing selection within immune genes that would bias the MK tests downward (Cagliani et al., 2012). These results show that viruses have been a major selective pressure in human evolution, and have had an influence on the rate of adaptation in the human proteome that extends far beyond antiviral defense proteins, and on a much greater scale than previously appreciated.

## Frequent adaptation in VIPs across mammals

The increased rate of adaptation in VIPs in the human lineage strongly suggests that VIPs in our dataset, 95% of which interact with modern viruses (Table S1), were also VIPs during past human evolution. It is also plausible that a substantial proportion of the VIPs we study are also VIPs in multiple mammalian lineages. Indeed, viruses infecting humans (including the ten viruses with the most VIPs) are known to have close viral relatives in many other mammals, with the exception of Hepatitis C Virus (HCV) for which only distant relatives are known and primarily in bats (Quan et al., 2013). There is also growing evidence that distantly related viruses tend to interact with overlapping sets of host proteins (Davis et al., 2015; Jager et al., 2012). We thus hypothesize that VIPs, while identified primarily in humans, may have also experienced frequent adaptation in mammals in general, with the possible exception of the VIPs interacting with HCV.

To test this hypothesis we use the Branch-Site Random Effect Likelihood test (BS-REL test) (Kosakovsky Pond et al., 2011) available in the HYPHY package (Pond et al., 2005) to detect episodes of adaptive evolution in each of the 44 branches of the mammalian tree used for the analysis (Methods). For a specific coding sequence, the BS-REL test estimates the proportion of codons where the rate of non-synonymous substitutions is higher than the rate of synonymous substitutions (dN/dS>1), which is a hallmark of adaptive evolution. The BS-REL test then compares two competing models of evolution, one with adaptive substitutions and one without adaptive substitutions, and decides which of the two models is the best fit. For each branch of the tree, the BS-REL test provides a *P*-value that corresponds to the probability that no adaptation occurred in the branch. The product of *P*-values across all branches in the tree then gives the probability that no adaptation occurred anywhere along the entire tree (Supplemental Methods).

The product of *P*-values is a good measure of whether a specific protein experienced adaptation in the history of mammalian evolution. In addition to presence/absence of adaptation, we assess the amount of adaptation experienced by a particular protein by estimating the average proportion of selected codons along all mammalian branches. For example we measure that on average the antiviral protein PKR has had 4.6% of positively selected codons across the 44 branches of the mammalian tree.

We compare the proportion of selected codons detected by the BS-REL test between VIPs and non-VIPs. The statistical power of the BS-REL test has been shown to depend strongly on the amount of constraint in a coding sequence, with higher constraint/purifying selection decreasing the ability to detect adaptation (Kosakovsky Pond et al., 2011). We confirm this in our dataset by observing a strong positive correlation between the pN/pS ratio in great apes and the proportion of selected codons across mammals estimated by the BS-REL test (Spearman’s rank correlation *ρ* = 0.34, *P*<2×10^−16^, n=9,861). We therefore use a purifying selection-wise permutation test that matches VIPs and non-VIPs with similar pN/pS ratios in order to compare VIPs and non-VIPs that experience similar levels of purifying selection and providing us with similar power to detect adaptation (Supplemental Methods and Figures S5 and S6).

The purifying selection-wise permutation test shows that adaptation has been much more common in VIPs than in non-VIPs across mammals (Figure 2). We estimate that VIPs have experienced 77% more adaptation compared to non-VIPs (Figure 2A). In total, this represents ~76,000 more adaptive amino acid changes in VIPs compared to non-VIPs. We further use an increasingly strict level of evidence for the presence of adaptation, by including only proteins with increasingly low products of P-values; that is, increasingly low probability that no adaptation occurred (Figure 2A). Figure 2A shows that VIPs with the strongest evidence of adaptation (product of *P*-values lower than 10^−9^) have a ~200% excess of strong signals of adaptation. This excess of adaptation in VIPs across mammals is due to i) more VIPs with signals of adaptation than non-VIPs, ii) more branches of the tree per VIP showing adaptation, and iii) a greater proportion of codons evolving adaptively per branch (Figure S7).

**Figure 2.**
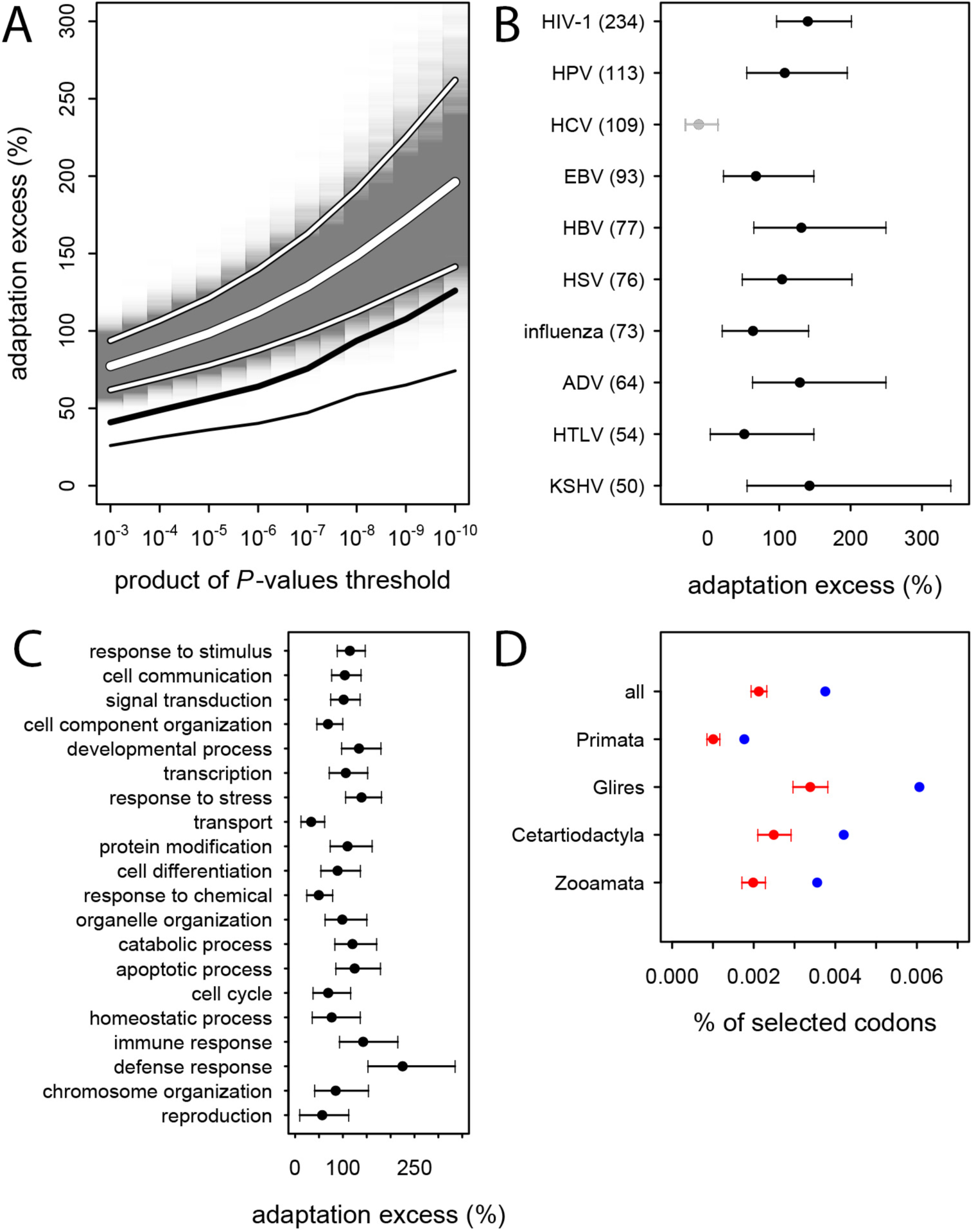
Excess of adaptation across mammals in VIPs. A) Thick white curve: average excess of adaptation in all VIPs. Narrow white curves: 95% confidence interval for the excess of adaptation in all VIPs. The grey background represents the density distributions of the excess of adaptation based on 5,000 iterations of the purifying selection-wise permutation test (Supplemental Methods). Thick black curve: excess of adaptation in non-immune VIPs. Narrow black curve: lower bound of the 95% confidence interval for the excess of adaptation in non-immune VIPs. B) Virus-by-virus excess of adaptation in VIPs. Black dot is the average excess and the represented interval is the 95% confidence interval. C) Excess of adaptation within the top 20 high level GO processes with the most VIPs. D) Proportions of selected codons in VIPs (blue dot) and non-VIPs (red dot and 95% confidence interval) in the mammalian clades represented by more than one species in the tree. All: entire tree. Primata: primates. Glires: rodents and rabbit. Cetartyodactyla: sheep, cow, pig. Zooamata: carnivores and horse. See also Tables S4 and S4 and Figures S5, S6, S7, S8 and S9.

As mentioned above, HCV stands out among the ten viruses with the largest number of VIPs in humans in that it has no known close viral relatives despite extensive screening of diverse mammalian species (Quan et al., 2013). If this reflects a true lack of close viral relatives of HCV, then we predict a limited excess of adaptation in HCV VIPs. In line with this prediction, the 109 VIPs of HCV are the only ones where we do not detect any excess of adaptation (Figure 2B) despite being one of the largest groups of VIPs. VIPs that interact with all other viruses all show substantial elevation of adaptation (Figure 2B).

GO processes with a strong excess of adaptation include cellular processes such as transcription, signal transduction, apoptosis, or post-translational protein modification, but also supracellular processes related to development (Figure 2C and Table S3). Importantly, VIPs with no known immune function (Table S4) show a 40% excess of adaptation, and a ~130% excess of adaptation in the proteins with the highest evidence of adaptation (Figure 2A). These results show that the arms race with viruses has strongly increased the rate of adaptation in a wide range of VIPs.

Since 95% of the VIPs were discovered for viruses infecting humans, it is possible that the observed excess of adaptation in VIPs in mammals is due to higher rates of adaptation exclusively in the primate branches of the mammalian tree (Figure S1). However, all mammalian clades in the tree show a similar excess of adaptation in VIPs (Figure 2D). Primates stand out due to their low overall proportions of positively selected codons compared to the other mammalian clades in the tree (Figure 2D). This is most likely due to a lower statistical power of the BS-REL test in the short primate branches (Kosakovsky Pond et al., 2011). In line with this, VIPs with strong signals of adaptation show such signals in all the mammalian clades represented (Figure 3). This includes well-known antiviral VIPs (Figure 3A), antiviral VIPs where adaptation was previously unknown (Figure 3B), and non-antiviral VIPs with diverse, well-studied functions in the mammalian hosts (Figure 3C). This phylogenetically widespread excess of adaptation implies that many of the VIPs annotated in humans were also VIPs for a substantial evolutionary time in a wide range of mammals.

**Figure 3.**
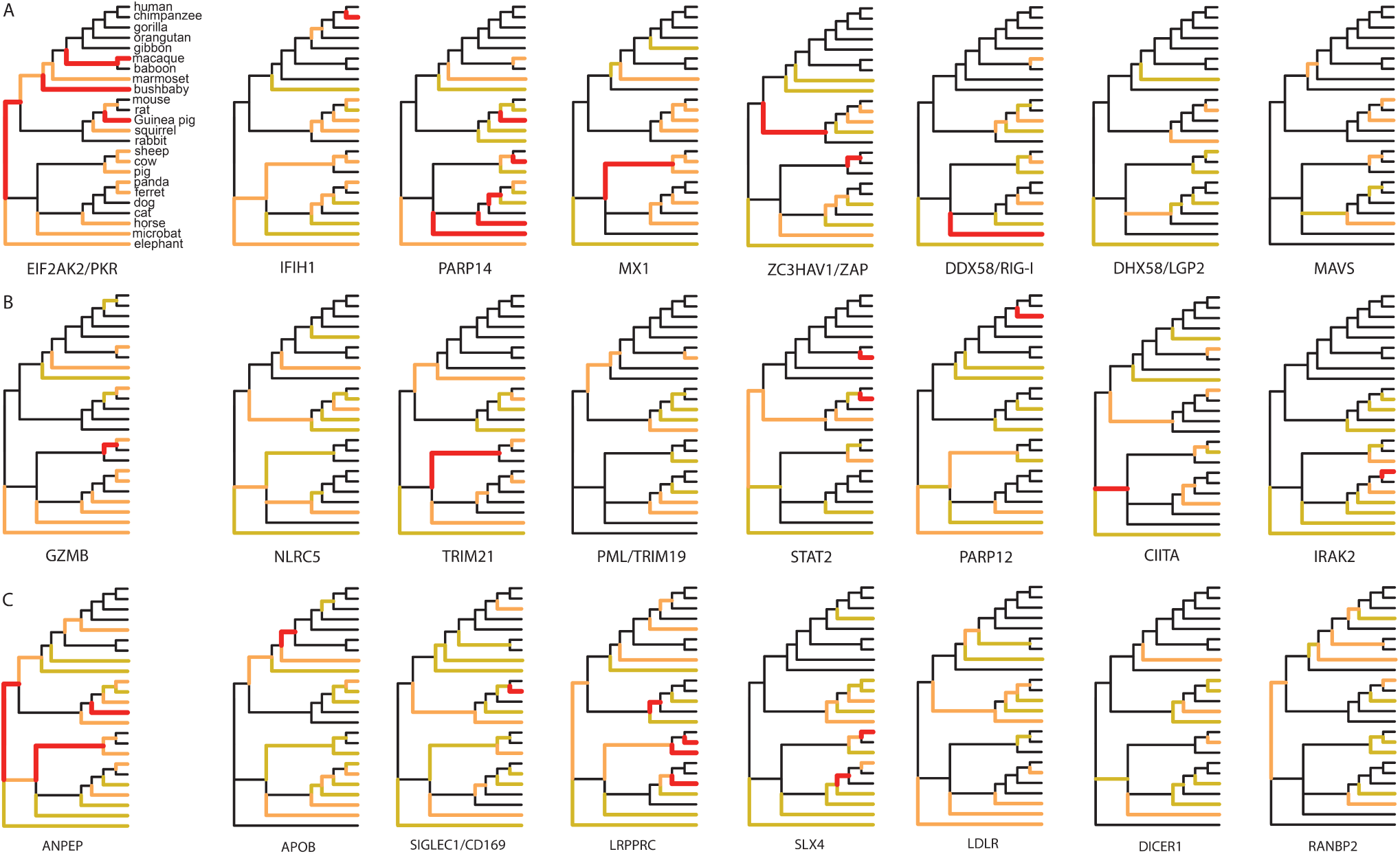
Examples of mammalian orthologs with adaptation spread across clades. A) Signals of adaptation in eight antiviral proteins with well-known adaptation. Red: BS-REL *P*≤0.001. Orange: BS-REL *P*≤0.05. Yellow: BS-REL *P*≤0.1. B) Top eight antiviral proteins with the highest number of branches under selection, and no previously know adaptive evolution. C) Top eight non-antiviral proteins with well-known functions and the highest number of branches under selection. Proteins are ordered according to the number of branches with signals of adaptation. All proteins in the figure have their product of BS-REL test *P*-values lower than 10^−9^.

## From global patterns of adaptation to understanding specific instances of adaptation to viruses: the case of coronaviruses and aminopeptidase N

We have shown that rates of adaptation are globally elevated in VIPs in humans and mammals in general, suggesting the existence of tens of thousands of isolated events of adaptations to a diverse range of viruses. Here, we test if our global approach has enough power to isolate new specific cases of adaptation to viruses by looking for instances where viruses are the plausible cause of adaptation in a VIP with no known antiviral activity. This is particularly relevant because, to our knowledge, the transferrin receptor is the only well understood case of a non-antiviral protein adapting in response to viruses (Demogines et al., 2013).

To identify a new non-antiviral protein we first exclude all VIPs with a well-known antiviral activity (Table S4) and then select all remaining VIPs with strong overall evidence of adaptation (Table S10) and at least 10 branches with signals of adaptation. Because we want to understand how adaptation to viruses proceeded, we then select proteins with i) at least one available tertiary structure, ii) amino acid level resolution of the interaction with one or more viruses, and iii) host tropism.

The most positively selected non-antiviral VIP that fulfills all these requirements is aminopeptidase N, abbreviated ANPEP, APN or CD13 (Mina-Osorio, 2008). The analysis of a phylogenetic tree including 84 mammals (Table S11) confirms pervasive adaptation of ANPEP across mammals, with 76 out of 165 branches in the tree evolving adaptively (Figure 4A). ANPEP is a cell-surface enzyme well known for its surprisingly wide range and diversity of functions (Mina-Osorio, 2008). It is used by group I coronaviruses as a receptor, including the Human Coronavirus 229E (HCoV-229E) (Yeager et al., 1992), Transmissible Gastroenteritis Virus (TGEV) (Delmas et al., 1992), Feline Coronavirus (FCoV) (Tresnan and Holmes, 1998), Canine Coronavirus (CCoV) (Tusell et al., 2007), Porcine Respiratory Coronavirus (PRCV) (Delmas et al., 1993) and Porcine Epidemic Diarrhea Virus (PEDV) (Oh et al., 2003). Reguera *et al.* (Reguera et al., 2012) have solved the structure of porcine ANPEP bound together with TGEV and PRCV. The authors identified in the extracellular domain of ANPEP 22 amino acids that form a surface of contact with TGEV and PRCV (Figure 4B) (Pettersen et al., 2004). The most important component of this contact surface for host tropism is a N-glycosylation site at position 736 in porcine ANPEP (orthologous position 739 in human ANPEP) that forms hydrogen bonds with TGEV and PRCV (Reguera et al., 2012; Tusell et al., 2007). Deleting this site abolishes the ability of TGEV and PRCV to bind porcine ANPEP (Reguera et al., 2012). Adding the glycosylation site in human ANPEP that natively lacks it transforms it into a receptor for TGEV and PRCV (Reguera et al., 2012).

**Figure 4.**
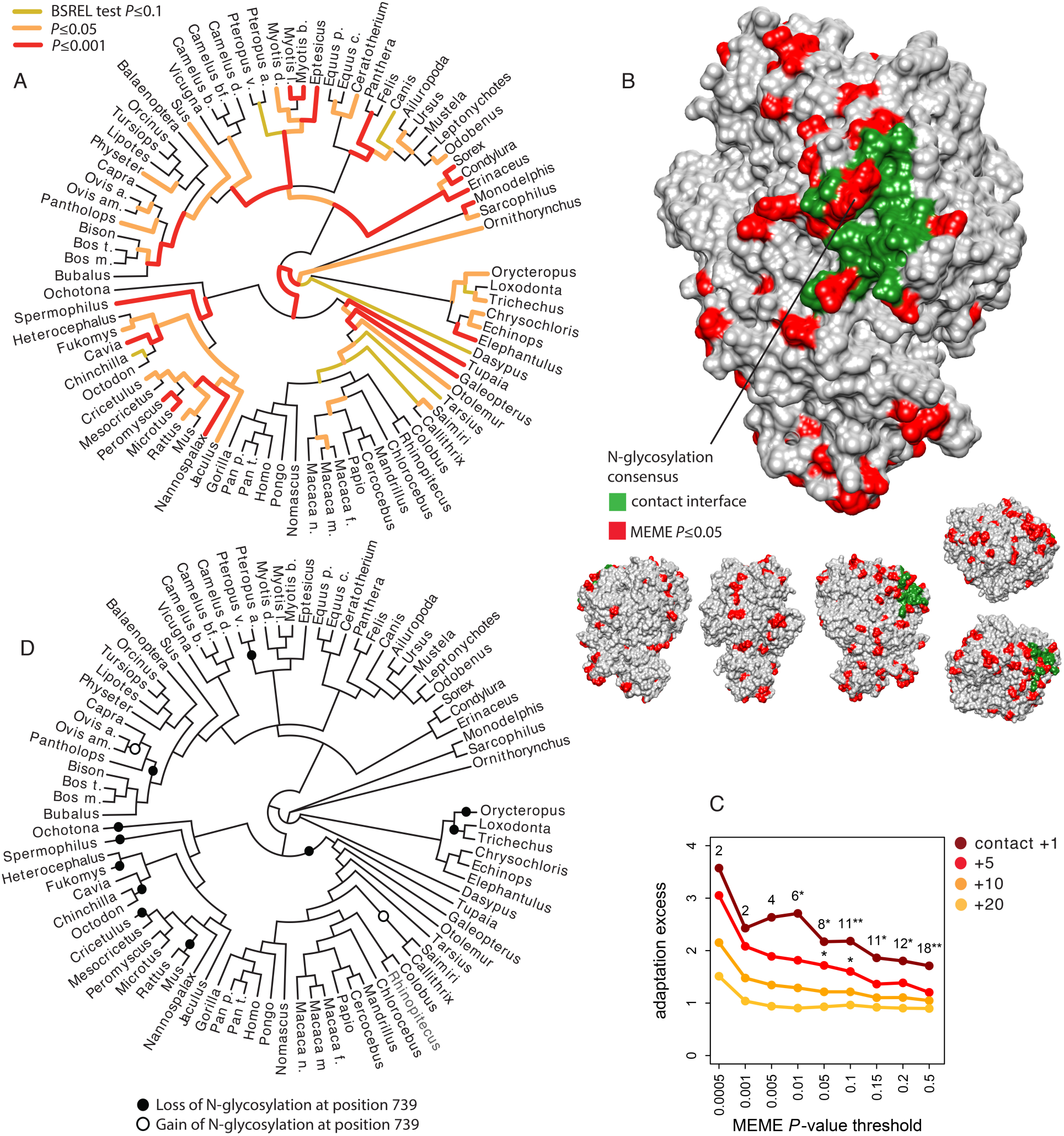
Patterns of adaptation to coronaviruses in aminopeptidase N. A) BS-REL test results for ANPEP in a tree of 84 mammalian species. Legend is on the figure. B) Contact surface with PRCV and TGEV on ANPEP structure (PDB 4FYQ). The figure includes visualizations of all the six different faces of ANPEP. Legend is in the figure. C) Excess of adaptation in and near the contact interface with PRCV and TGEV. Within the contact interface plus a given number of neighboring amino acids (one, five, ten or 20 in the figure), adaptation excess (y axis) is defined as the number of observed codons with a MEME P-value lower than the P-value threshold on the x axis, divided by the average number of codons under the same P-value threshold obtained after randomizing the location of adaptation signals over the entire ANPEP coding sequence 5,000 times. Dark red curve: adaptation excess within the contact interface with TGEV and PRCV plus one neighboring amino acid. Red curve: plus five neighboring amino acids. Orange: plus ten neighboring amino acids. Light orange: plus 20 neighboring amino acids. Numbers in the figure represent the number of adapting codons, and the stars give the significance of the excess. One star: excess *P*≤0.05. Two stars: *P*≤0.01. D) Losses and gains of the N-glycosylation across the mammalian phylogeny. See also Tables S10 and S11.

We use the MEME test from HYPHY (Murrell et al., 2012) to identify codons in ANPEP that were under episodic adaptive evolution in mammals. MEME detects significant adaptation (MEME *P*≤0.05) in 85 of the 931 aligned codons. Interestingly, several of these adaptively evolving codons are within, or right next to the surface of contact with TGEV and PRCV (Figure 4B,C). The codons in contact with TGEV and PRCV and their neighbors are strongly enriched in adaptation compared to ANPEP codons as a whole (Figure 4C). This enrichment fades very rapidly as one gets further from the surface of contact with TGEV and PRCV, which shows that it is due to the interaction with viruses, and not to a more diffuse, less specific enrichment within a wider segment of ANPEP (Figure 4C).

Adaptively evolving codons in the contact surface with TGEV and PRCV most notably include two codons within the consensus motif for the N-glycosylation site responsible for host tropism (Figure 4B). N-glycosylation is governed by a three amino acids consensus, NX[ST], where X can be anything except proline (Bause, 1983). The first and third positions in the consensus evolved adaptively in mammals (MEME *P*=0.005 for both). The ancestral states of the two positions shows that the mammalian ancestor had a fully functional consensus for N-glycosylation, and that the consensus was lost independently 11 times in mammals, either by modification of the first or by modification of the third position (Figure 4D). The consensus was regained only two times after loss (Figure 4D). This means that the signals of adaptation detected at the first and third positions in the consensus mainly reflect parallel, adaptive losses of the N-glycosylation site in multiple mammalian lineages. Given the crucial role of this N-glycosylation site in the binding of TGEV, PRCV, FCoV and CCoV to ANPEP, it is very likely that these parallel adaptive losses were the result of the selective pressure exerted by ancient coronaviruses.

## Discussion

Here, we have shown that viruses have been a major selective pressure in the evolution of the mammalian proteome. Indeed viruses appear to drive ~30% of all adaptive amino acid changes in the conserved part of the human proteome, as evidenced by the MK test. Furthermore, the footprints of the arms race with viruses are visible in a large number of VIPs, and in a broad range of mammals. Importantly, we find a substantial enrichment in strong signals of adaptation in VIPs with no known antiviral or other immune functions (Figures 1F and 2A). Instead adaptation to viruses is visible in VIPs with a very diverse range of functions including such core functions as transcription or signal transduction (Figure 2C and Table S3). Our results thus draw a broader picture where the war against viruses is a global war that involves not only the specialized soldiers of the antiviral response, but also the entire population of host proteins that come into contact with viruses. The best-known case of a housekeeping protein having adapted in response to viruses, the transferrin receptor, may in this respect represent the rule more than an exception (Demogines et al., 2013).

Although we already find a strong signal of increased adaptation, the amount of adaptive evolution that can be attributed to viruses is probably underestimated by our analysis. First, there may still be many undiscovered VIPs. Within the past few years, there has been no sign that the pace of discovery of new VIPs is slowing down (Figure S2). A slowdown would happen if we were getting close to the point where only few, hard to discover VIPs remain to be found. This means that a substantial number of proteins classified as non-VIPs in this analysis actually are VIPs. This makes the current non-VIPs a conservative control.

Second, adaptation in response to viruses is most likely not restricted to proteins that physically interact with viruses. For example, adaptation to viruses might happen in proteins that act downstream of VIPs in signaling cascades, or in non-coding sequences that regulate the expression of VIPs.

Third, not all of the 1,256 VIPs we use here have been consistently interacting with viruses during evolution. Most VIPs in the dataset (95%) were discovered in humans, and how frequently these VIPs have also been interacting with viruses in other mammals is currently unknown. Some VIPs like PKR have probably been in very frequent contact with diverse viruses. Conversely, other VIPs may have been in contact with viruses for a very limited evolutionary time in mammals, and only in a limited range of lineages. This would apply to VIPs that interact with viruses with a limited host range and few other phylogenetically closely related viruses, as is the case with HCV (Figure 2B).

Our analysis greatly extends the prospects for paleovirology by making it possible to detect past infections with great statistical power (Emerman and Malik, 2010; Patel et al., 2011). Indeed, globally increased adaptation in VIPs means that we have far more statistical power to study the adaptive footprints viruses have left in host proteins. This is a very complementary approach to the usual scans for integrated viral genomes in host genomes, especially for viruses that do not infect the germ line or for viruses that integrate in the host genome only rarely (Katzourakis and Gifford, 2010). In the dataset analyzed here, only HCV does not show any excess of adaptation in mammals. Two of the ten viruses listed in Figure 2B, HIV-1 and HBV, have VIPs with highly increased adaptation in humans. In other words, these results demonstrate that viruses related to HIV-1 and HBV have repeatedly infected humans since their divergence with chimpanzees. These results show that it should be possible to detect the footprint of past infections by specific viruses in specific evolutionary lineages with great statistical power.

Globally increased adaptation in VIPs also raises the question of pleiotropy. To what extent have adaptive amino acid changes in VIPs affected their native functions in the host? We show that there has been so much adaptation in VIPs that it is very hard to imagine that none of these adaptive events had any consequences on host phenotypes. Interestingly, VIPs tend to be multifunctional proteins. Indeed they represent 13% of all the orthologs in the analysis, 33% of the orthologs with 60 or more annotated GO processes, and 40% of orthologs with 100 or more GO processes (Figure S8A). Pleiotropy is more likely in proteins with many functions (He and Zhang, 2006), and the subset of VIPs with many annotated GO processes has an excess of adaptation that is very similar to the one observed when using all VIPs (Figure S8B). Adaptation to viruses could thus have affected the evolution of host phenotypes in unexpected ways. In this respect, it is particularly intriguing that VIPs have experienced highly increased rates of adaptation within host functions such as development or neurogenesis (Table S3).

Finally, our results might explain a puzzling observation that rates of protein adaptation appear relatively invariant across different biological functions (Bierne and Eyre-Walker, 2004). Because viruses (and likely other pathogens) interact with diverse proteins across most GO functions, they elevate the rate of adaptation across the whole proteome in a way that appears independent of specific functions in the GO analysis. We argue that grouping genes together based on the way they interact with diverse pathogens or other environmental stimuli might be a profitable way for discerning the nature of selective pressures that have molded animal genomes.

In conclusion, our analysis demonstrates that viruses have exerted a very powerful selective pressure across the breadth of the mammalian proteome, and suggests the possibility that pathogens in general are the key driver of protein adaptation in mammals and likely other lineages and might have driven many pleiotropic effects on diverse biological functions.

## Methods

### Manual curation of virus-interacting proteins

We identified 1,256 VIPs out of a total of 9,861 proteins with orthologs in the genomes of the 24 mammals included in the analysis (Figure S1 and Tables S1 and S2). Annotation of the 1,256 VIPs was performed by querying PUBMED for publications reporting interactions found by low-throughput methods (Gray et al., 2015) (http://www.ncbi.nlm.nih.gov/pubmed). We completed our own annotations of 982 VIPs with 274 additional proteins identified with low throughput methods listed in the VirHostNet (http://virhostnet.prabi.fr/) (Guirimand et al., 2015) and HPIDB (http://www.agbase.msstate.edu/hpi/main.html) (Kumar and Nanduri, 2010) databases. See Supplemental Methods for details.

### Multiple alignments of mammalian orthologs

We aligned 9,861 orthologous Ensembl coding sequences (Flicek et al., 2012) from 24 mammals using a combination of Blat (Kent, 2002) and PRANK (Fletcher and Yang, 2010; Jordan and Goldman, 2012; Loytynoja and Goldman, 2008). See Supplemental Methods for details.

### Annotation of antiviral and immune mammalian orthologs

In addition to annotating VIPs, we also annotated which VIPs have known antiviral or broader immune activities. See Supplemental Methods for details.

### Quantifying adaptation in human

We use the classic and the asymptotic McDonald-Kreitman tests to compare adaptation in VIPs and non-VIPs in human. See Supplemental Methods for details.

### Quantifying adaptation in the mammalian phylogeny

We use the CDS alignments of the 24 mammals to quantify adaptation in their phylogeny. To do so we use the Branch-Site REL test (Kosakovsky Pond et al., 2011) from the HYPHY package (Pond et al., 2005). See Supplemental Methods for details.

### The pN/pS-based purifying selection-wise permutation test

We created a permutation test that compares VIPs and non-VIPs with the same amount of purifying selection. This is achieved by using the pN/pS ratio as a proxy for purifying selection. See Supplemental Methods for details.

### Retrieving of ANPEP mammalian coding sequences

We analyzed patterns of adaptation in ANPEP in a tree of mammals including 84 species. These species are the ones with annotated, known or predicted mRNAs (Table S11 for their Genbank identifiers). The coding sequences were extracted from the mRNAs and aligned with PRANK.

### Gene Ontology-matching control samples

We created a permutation scheme that compares VIPs with random samples of non-VIPs with the same representation of GO categories. See Supplemental Methods for details.

## Author Contributions

DE and DAP conceived and designed the experiments. DE, LC and CG performed the experiments. DE and DAP interpreted the results. DE and DAP wrote the paper.

## Acknowledgements

We thank Kerry Samerotte, Sandeep Venkataram, Emily Ebel and Pleuni Pennings for comments on the manuscript. This work is funded by NIH grants R01GM089926 and R01GM097415 to DAP.

## References

(2015). Gene Ontology Consortium: going forward. Nucleic Acids Res 43, D1049–1056.

Abecasis, G.R., Auton, A., Brooks, L.D., DePristo, M.A., Durbin, R.M., Handsaker, R.E., Kang, H.M., Marth, G.T., and McVean, G.A. (2012). An integrated map of genetic variation from 1,092 human genomes. Nature 491, 56–65.

Ashburner, M., Ball, C.A., Blake, J.A., Botstein, D., Butler, H., Cherry, J.M., Davis, A.P., Dolinski, K., Dwight, S.S., Eppig, J.T., et al. (2000). Gene ontology: tool for the unification of biology. The Gene Ontology Consortium. Nat Genet 25, 25–29.

Bause, E. (1983). Structural requirements of N-glycosylation of proteins. Studies with proline peptides as conformational probes. Biochem J 209, 331–336.

Bierne, N., and Eyre-Walker, A. (2004). The genomic rate of adaptive amino acid substitution in Drosophila. Mol Biol Evol 21, 1350–1360.

Cagliani, R., Guerini, F.R., Fumagalli, M., Riva, S., Agliardi, C., Galimberti, D., Pozzoli, U., Goris, A., Dubois, B., Fenoglio, C., et al. (2012). A trans-specific polymorphism in ZC3HAV1 is maintained by long-standing balancing selection and may confer susceptibility to multiple sclerosis. Mol Biol Evol 29, 1599–1613.

Cagliani, R., Riva, S., Fumagalli, M., Biasin, M., Caputo, S.L., Mazzotta, F., Piacentini, L., Pozzoli, U., Bresolin, N., Clerici, M., et al. (2011). A positively selected APOBEC3H haplotype is associated with natural resistance to HIV-1 infection. Evolution 65, 3311–3322.

Charlesworth, J., and Eyre-Walker, A. (2008). The McDonald-Kreitman test and slightly deleterious mutations. Mol Biol Evol 25, 1007–1015.

Davis, Z.H., Verschueren, E., Jang, G.M., Kleffman, K., Johnson, J.R., Park, J., Von Dollen, J., Maher, M.C., Johnson, T., Newton, W., et al. (2015). Global mapping of herpesvirus-host protein complexes reveals a transcription strategy for late genes. Mol Cell 57, 349–360.

Delmas, B., Gelfi, J., L’Haridon, R., Vogel, L.K., Sjostrom, H., Noren, O., and Laude, H. (1992). Aminopeptidase N is a major receptor for the entero-pathogenic coronavirus TGEV. Nature 357, 417–420.

Delmas, B., Gelfi, J., Sjostrom, H., Noren, O., and Laude, H. (1993). Further characterization of aminopeptidase-N as a receptor for coronaviruses. Adv Exp Med Biol 342, 293–298.

Demogines, A., Abraham, J., Choe, H., Farzan, M., and Sawyer, S.L. (2013). Dual host-virus arms races shape an essential housekeeping protein. PLoS Biol 11, e1001571.

Dyer, M.D., Murali, T.M., and Sobral, B.W. (2008). The landscape of human proteins interacting with viruses and other pathogens. PLoS Pathog 4, e32.

Elde, N.C., Child, S.J., Geballe, A.P., and Malik, H.S. (2009). Protein kinase R reveals an evolutionary model for defeating viral mimicry. Nature 457, 485–489.

Emerman, M., and Malik, H.S. (2010). Paleovirology--modern consequences of ancient viruses. PLoS Biol 8, e1000301.

Eyre-Walker, A., and Keightley, P.D. (2009). Estimating the rate of adaptive molecular evolution in the presence of slightly deleterious mutations and population size change. Mol Biol Evol 26, 2097–2108.

Fletcher, W., and Yang, Z. (2010). The effect of insertions, deletions, and alignment errors on the branch-site test of positive selection. Mol Biol Evol 27, 2257–2267.

Flicek, P., Amode, M.R., Barrell, D., Beal, K., Brent, S., Carvalho-Silva, D., Clapham, P., Coates, G., Fairley, S., Fitzgerald, S., et al. (2012). Ensembl 2012. Nucleic Acids Res 40, D84–90.

Fumagalli, M., Cagliani, R., Riva, S., Pozzoli, U., Biasin, M., Piacentini, L., Comi, G.P., Bresolin, N., Clerici, M., and Sironi, M. (2010). Population genetics of IFIH1: ancient population structure, local selection, and implications for susceptibility to type 1 diabetes. Mol Biol Evol 27, 2555–2566.

Gray, K.A., Yates, B., Seal, R.L., Wright, M.W., and Bruford, E.A. (2015). Genenames.org: the HGNC resources in 2015. Nucleic Acids Res 43, D1079–1085.

Guirimand, T., Delmotte, S., and Navratil, V. (2015). VirHostNet 2.0: surfing on the web of virus/host molecular interactions data. Nucleic Acids Res 43, D583–587.

Halehalli, R.R., and Nagarajaram, H.A. (2015). Molecular principles of human virus protein-protein interactions. Bioinformatics 31, 1025–1033.

He, X., and Zhang, J. (2006). Toward a molecular understanding of pleiotropy. Genetics 173, 1885–1891.

Jager, S., Cimermancic, P., Gulbahce, N., Johnson, J.R., McGovern, K.E., Clarke, S.C., Shales, M., Mercenne, G., Pache, L., Li, K., et al. (2012). Global landscape of HIV-human protein complexes. Nature 481, 365–370.

Jordan, G., and Goldman, N. (2012). The effects of alignment error and alignment filtering on the sitewise detection of positive selection. Mol Biol Evol 29, 1125–1139.

Kaelber, J.T., Demogines, A., Harbison, C.E., Allison, A.B., Goodman, L.B., Ortega, A.N., Sawyer, S.L., and Parrish, C.R. (2012). Evolutionary reconstructions of the transferrin receptor of Caniforms supports canine parvovirus being a re-emerged and not a novel pathogen in dogs. PLoS Pathog 8, e1002666.

Katzourakis, A., and Gifford, R.J. (2010). Endogenous viral elements in animal genomes. PLoS Genet 6, e1001191.

Keightley, P.D., and Eyre-Walker, A. (2007). Joint inference of the distribution of fitness effects of deleterious mutations and population demography based on nucleotide polymorphism frequencies. Genetics 177, 2251–2261.

Kent, W.J. (2002). BLAT--the BLAST-like alignment tool. Genome Res 12, 656–664.

Kerns, J.A., Emerman, M., and Malik, H.S. (2008). Positive selection and increased antiviral activity associated with the PARP-containing isoform of human zinc-finger antiviral protein. PLoS Genet 4, e21.

Kimura, M. (1977). Preponderance of synonymous changes as evidence for the neutral theory of molecular evolution. Nature 267, 275–276.

Kosakovsky Pond, S.L., Murrell, B., Fourment, M., Frost, S.D., Delport, W., and Scheffler, K. (2011). A random effects branch-site model for detecting episodic diversifying selection. Mol Biol Evol 28, 3033–3043.

Kumar, R., and Nanduri, B. (2010). HPIDB--a unified resource for host-pathogen interactions. BMC Bioinformatics 11 Suppl 6, S16.

Liu, H.L., Wang, Y.Q., Liao, C.H., Kuang, Y.Q., Zheng, Y.T., and Su, B. (2005). Adaptive evolution of primate TRIM5alpha, a gene restricting HIV-1 infection. Gene 362, 109–116.

Loytynoja, A., and Goldman, N. (2008). A model of evolution and structure for multiple sequence alignment. Philos Trans R Soc Lond B Biol Sci 363, 3913–3919.

McDonald, J.H., and Kreitman, M. (1991). Adaptive protein evolution at the Adh locus in Drosophila. Nature 351, 652–654.

Mellacheruvu, D., Wright, Z., Couzens, A.L., Lambert, J.P., St-Denis, N.A., Li, T., Miteva, Y.V., Hauri, S., Sardiu, M.E., Low, T.Y., et al. (2013). The CRAPome: a contaminant repository for affinity purification-mass spectrometry data. Nat Methods 10, 730–736.

Messer, P.W. (2013). SLiM: simulating evolution with selection and linkage. Genetics 194, 1037–1039.

Messer, P.W., and Petrov, D.A. (2013). Frequent adaptation and the McDonald-Kreitman test. Proc Natl Acad Sci U S A 110, 8615–8620.

Mina-Osorio, P. (2008). The moonlighting enzyme CD13: old and new functions to target. Trends Mol Med 14, 361–371.

Murrell, B., Wertheim, J.O., Moola, S., Weighill, T., Scheffler, K., and Kosakovsky Pond, S.L. (2012). Detecting individual sites subject to episodic diversifying selection. PLoS Genet 8, e1002764.

Oh, J.S., Song, D.S., and Park, B.K. (2003). Identification of a putative cellular receptor 150 kDa polypeptide for porcine epidemic diarrhea virus in porcine enterocytes. J Vet Sci 4, 269–275.

Patel, M.R., Emerman, M., and Malik, H.S. (2011). Paleovirology - ghosts and gifts of viruses past. Curr Opin Virol 1, 304–309.

Pettersen, E.F., Goddard, T.D., Huang, C.C., Couch, G.S., Greenblatt, D.M., Meng, E.C., and Ferrin, T.E. (2004). UCSF Chimera--a visualization system for exploratory research and analysis. J Comput Chem 25, 1605–1612.

Pond, S.L., Frost, S.D., and Muse, S.V. (2005). HyPhy: hypothesis testing using phylogenies. Bioinformatics 21, 676–679.

Prado-Martinez, J., Sudmant, P.H., Kidd, J.M., Li, H., Kelley, J.L., Lorente-Galdos, B., Veeramah, K.R., Woerner, A.E., O’Connor, T.D., Santpere, G., et al. (2013). Great ape genetic diversity and population history. Nature 499, 471–475.

Quan, P.L., Firth, C., Conte, J.M., Williams, S.H., Zambrana-Torrelio, C.M., Anthony, S.J., Ellison, J.A., Gilbert, A.T., Kuzmin, I.V., Niezgoda, M., et al. (2013). Bats are a major natural reservoir for hepaciviruses and pegiviruses. Proc Natl Acad Sci U S A 110, 8194–8199.

Reguera, J., Santiago, C., Mudgal, G., Ordono, D., Enjuanes, L., and Casasnovas, J.M. (2012). Structural bases of coronavirus attachment to host aminopeptidase N and its inhibition by neutralizing antibodies. PLoS Pathog 8, e1002859.

Sawyer, S.L., Emerman, M., and Malik, H.S. (2004). Ancient adaptive evolution of the primate antiviral DNA-editing enzyme APOBEC3G. PLoS Biol 2, E275.

Sawyer, S.L., Emerman, M., and Malik, H.S. (2007). Discordant evolution of the adjacent antiretroviral genes TRIM22 and TRIM5 in mammals. PLoS Pathog 3, e197.

Sawyer, S.L., Wu, L.I., Emerman, M., and Malik, H.S. (2005). Positive selection of primate TRIM5alpha identifies a critical species-specific retroviral restriction domain. Proc Natl Acad Sci U S A 102, 2832–2837.

Sironi, M., Biasin, M., Cagliani, R., Forni, D., De Luca, M., Saulle, I., Lo Caputo, S., Mazzotta, F., Macias, J., Pineda, J.A., et al. (2012). A common polymorphism in TLR3 confers natural resistance to HIV-1 infection. J Immunol 188, 818–823.

Tresnan, D.B., and Holmes, K.V. (1998). Feline aminopeptidase N is a receptor for all group I coronaviruses. Adv Exp Med Biol 440, 69–75.

Tusell, S.M., Schittone, S.A., and Holmes, K.V. (2007). Mutational analysis of aminopeptidase N, a receptor for several group 1 coronaviruses, identifies key determinants of viral host range. J Virol 81, 1261–1273.

Vasseur, E., Patin, E., Laval, G., Pajon, S., Fornarino, S., Crouau-Roy, B., and Quintana-Murci, L. (2011). The selective footprints of viral pressures at the human RIG-I-like receptor family. Hum Mol Genet 20, 4462–4474.

Yeager, C.L., Ashmun, R.A., Williams, R.K., Cardellichio, C.B., Shapiro, L.H., Look, A.T., and Holmes, K.V. (1992). Human aminopeptidase N is a receptor for human coronavirus 229E. Nature 357, 420–422.

